# Nonlinear collision between anisotropic propagating waves in mouse somatosensory cortex

**DOI:** 10.1101/2021.01.04.425215

**Authors:** M. Di Volo, I. Férézou

## Abstract

How does cellular organization shape the spatio-temporal patterns of activity in the cortex while processing sensory information? After measuring the propagation of activity in the mouse primary somatosensory cortex (S1) in response to single whisker deflections with Voltage Sensitive Dye (VSD) imaging, we developed a two dimensional mean field model of S1. We observed that, for strong enough excitatory cortical interactions, whisker deflections generate a propagating wave in S1. We developed an inversion method that reconstructs model parameters from VSD data, revealing that a spatially heterogeneous organization of synaptic strengths between pyramidal neurons in S1 is likely to be responsible for the anisotropic spatio-temporal patterns of activity measured experimentally. Finally, we report that two consecutive stimuli activating different spatial locations in S1 generate two waves which collide sub-linearly. In the model, such sub-linear interaction is explained by a lower sensitivity to external perturbations of neural networks during activated states.

## Introduction

The 4th layer of the rodent primary somatosensory (S1) cortex has a remarkable cellular organization, with obvious structures, named “barrels”, laid out with a specific topology corresponding to the whiskers on the snout of the animal (Woolsey & Van der Loos, 1970). A precise tactile stimulation of a specific whisker on the snout of the animal activates primarily the corresponding barrel in S1 (Estebanez et al., 2018; Feldmeyer et al., 2013; Petersen, 2007). Because of its structure, considered as a manifestation of the columnar functional organization of the cerebral cortex, S1 has been largely studied in order to understand the way cortex processes tactile sensory information. It constitutes an ideal framework where to study the emergence of complex spatio-temporal patterns of activity in response to discrete sensory stimulations. Indeed, while a temporally defined individual whisker stimulation activates at first its corresponding barrel in S1, the activity rapidly propagates to the neighboring barrel-related columns, activating in a few tens of milliseconds almost the whole S1. Such spatio-temporal propagation can be measured in S1 using VSD imaging (Ferezou et al., 2006, 2007; Petersen et al., 2003), revealing that the simple stimulation of one whisker generates a complex representation across space and time in S1. In this work we investigate, by combining VSD imaging (VSDi) and computational modeling, how spatially distributed cellular connectivity in S1 regulates the emergence of anisotropic propagating waves in S1. Moreover, we characterize S1 response to more complex stimuli (more precisely, successive deflection of two distinct whiskers), where the collision between propagating waves of activity (originating at different locations) determines the way tactile stimulations are represented in the cortex.

Measuring S1 activity in response to tactile stimulations with VSD, we developed a biologically realistic two dimensional cortical model of S1 composed by interconnected networks of excitatory and inhibitory neurons. The model we propose employs a mean field approach to mimic the activity of populations of excitatory and inhibitory neurons. This approach has the advantage to describe the dynamics of a population of neurons with a low dimensional model (Boustani & Destexhe, 2009; Wilson & Cowan, 1972) but, at the same time, to be capable to encode biologically realistic ingredients (indeed we have recently shown that it is applicable to any spiking neuronal model, (Carlu et al., 2020)). A previous version of this model (one dimensional in space) has been employed to study the emergence of propagating waves in the primary visual cortex of the awake monkey (Chemla et al., 2019; Zerlaut et al., 2018). For a quantitative comparison with VSD images, we have here developed a two dimensional model that also includes spike frequency adaptation observed in excitatory regular spiking neurons in the cortex (Destexhe, 2009; Di Volo et al., 2019). Moreover, we have employed the model to predict the activity in S1 in response to the successive stimulation of two different but spatially close whiskers. The stimulation of one of the two whiskers produces a propagating wave that in few milliseconds reaches the spatial location of the representation of the other whisker. As a result, if the two stimuli are presented at a close time interval the two propagating waves collide. The model predicts a sublinear interaction between propagating waves in S1 due to lower sensitivity to external stimuli of cortical populations during high levels of activity. This prediction has been directly tested in experiments where we have indeed measured a suppressive wave (sublinear interaction) that propagates from the second stimuli location towards the first one. A similar observation has been recently reported in the primary visual cortex of the monkey where two consecutive and spatially close visual stimuli cause the collision of propagating waves with a suppressive interaction (Chemla et al., 2019). Our results reveal that this phenomenon is generally present in different animal models and brain regions.

## Results

### Anisotropic structure of propagating waves in S1

We measured the spatio-temporal activity of the mouse S1 at 500 Hz by means of VSD imaging (Grinvald & Hildesheim, 2004; Orbach et al., 1985). As previously reported (Ferezou et al., 2006) a small deflection of the right C2 whisker (see methods), evokes a response emerging in less than ten milliseconds at the corresponding barrel-related column location in the left S1. The signal then propagates in space during the following milliseconds, as it can be observed by looking at the pattern of activity at different time frames (Fig. 1A-B). This propagation is not uniform, as further revealed by linescan plots (time on x axis and space on y axis) of the VSD signal from lines drawn in different directions (Fig. 1C). Indeed, according to their orientation, the linescan plots show different spatio-temporal patterns of activity that are not symmetric with respect to the initial activation site. Dashed lines in Fig. 1C show the estimation of propagation velocity in different directions. The velocity is estimated by calculating the spatial location when the VSD signal exceeds a fixed threshold. In contrast to the intensity of the VSD signal, we found that the velocity of propagation (v_c = 0.1mm/ms) is approximately the same in all directions. In next sections we develop a mean field model of S1 to study how the emergent anisotropic structure of VSD activity depends on an heterogeneous distribution of neuronal connectivity in space.

**Figure 1:**
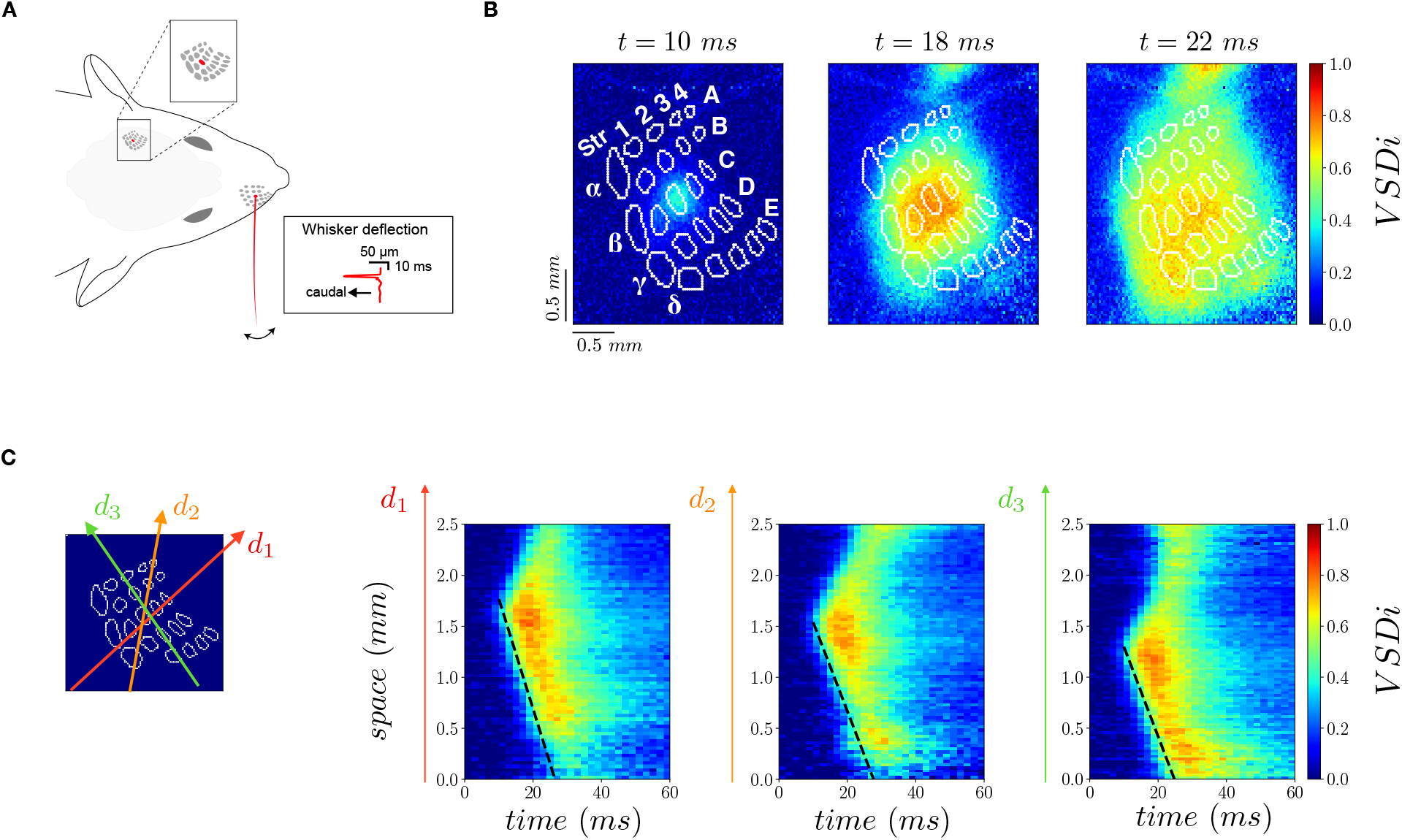
Experimental protocol and anisotropic propagating waves in S1. **A:** Experimental setup where the S1 is imaged at 500 images per second while the C2 whisker is stimulated with a piezoelectric actuator (2.7° displacement, with a 2-ms rising ramp, 2-ms plateau, and 2-ms fall). **B:** Color plots show images at different time frames after the beginning of the C2 whisker deflection (t = 0 ms). Intensity of the VSD signal is color-coded and normalized to the maximum intensity in space and time (average of n = 40 trials). The layer 4 barrel map reconstructed from a post hoc cytochrome oxidase staining is shown as overlaid white dots. **C:** The three arrows (d1. red, d2. orange and d3. green) indicate different spatial directions that we employ for a spatio-temporal visualization of VSD activity (linescan plots on the right). We have estimated, in each direction, the velocity of propagation through a fitting procedure based on the rise of activity in time at each spatial location. The fits (v_c_ = 0.1 +/- 0.01 m/s for the three directions) are reported as dashed black lines on the plots.

### A computational model for propagating waves in sensory cortex

We developed a two dimensional computational model of S1 to be compared with imaging data, by extending a mean-field model recently employed to reproduce accurately, in one dimension, VSD signals evoked by visual stimuli in the primary visual cortex of awake primates (Chemla et al., 2019; Zerlaut et al., 2018). We thus performed a discretization in space where each node of the resulting lattice is composed by two populations of interacting neurons, excitatory regular-spiking (RS) neurons, and inhibitory fast-spiking (FS) neurons (Fig. 2A). Excitatory RS neurons, in contrast to inhibitory FS cells, are characterized by spike frequency adaptation, in agreement with experimental observations (Destexhe, 2009). The mean field model gives access to the average membrane potential of the whole population, from which we estimate VSD intensity (see methods for details). A remarkable propriety of propagating waves in S1 is their spatial extent. Indeed the stimulation, as well as the resulting first activation in S1, is extremely confined spatially (Fig. 1 A). Nevertheless, the wave propagates far away from the activation site at speed vc with high intensity. The intensity of the signal in space is approximately the same of the intensity at the source (see Fig.1C, direction d3). A fundamental ingredient for stimulus-evoked propagating waves at the mesoscopic scale is the conduction velocity of the axons in superficial layers (Muller et al., 2018). Moreover, stimulus-evoked activity propagates with sustained intensity in bistable systems where the inputs traveling along axons trigger, with a space-dependent time delay, jumps from a low to a higher activity state (Capone et al., 2019). We employ, for each population of neurons (RS and FS), a recently developed mean field model with spike frequency adaptation (Di Volo et al., 2019). Spike frequency adaptation can give rise to a bidirectional switch from low to high activity states (Compte et al., 2003; Jercog et al., 2017; Mattia & Sanchez-Vives, 2012). Indeed, a pulse of external excitatory drive to the network can provoke the jump from a low to a high activity state. Because spike frequency adaptation builds up with increasing excitatory neurons activity, the activity then subsequently decreases. The hyperpolarization of excitatory neurons leads then to a decrease of activity and a jump back to the low activity state (Di Volo et al., 2019). When arranging several of these populations in a homogeneous space, a local input can potentially trigger a chain of jumps to high activity states that spreads in space. Nevertheless, the intensity of such activity is crucially shaped by heterogeneities in the system. Indeed, in a large variety of fields where propagating waves have been under scrutiny, e.g. experiments on the Belousov-Zhabotinsky reaction (BZR) (Steinbock et al., 1995) or on cardiac tissue (Bub et al., 2002), it has been shown that spatial heterogeneity can strongly affect the emergent structure of their spatiotemporal-dynamics. In our model, a crucial role is played by the spatial distribution of coupling strength between subpopulations of neurons. In this work we make the hypothesis that the coupling strength from a node (x_1_, y_1_) to another node (x,y) in space (Fig. 2A), namely 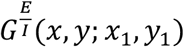, is exponentially decreasing with the Euclidean distance *d*. Moreover, we assume this coupling to be proportional to the strength of incoming synapses in (x,y), say 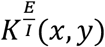 (see methods). The higher 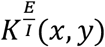, the stronger is the excitatory/inhibitory (E/I) input incoming in one neuron in (x,y). In our mean field formalism, *K^E^*(*x,y*) can be interpreted as the average amount of incoming excitatory connections in (x,y), or as the average strength of excitatory synaptic conductances for one neuron in (x,y). Notice that FS neurons do not have spike frequency adaptation and, as a result, they have a higher gain with respect to RS neurons (Destexhe, 2009). In the case where *K^E^*(*x, y*) = *K^I^*(*x, y*) = *K* is extracted from a spatially uniform distribution around 1 (see Fig. 2B), we observe that a pulse of excitatory activity in the location of C2 activates a propagating wave that spreads uniformly in space with a speed equal to the axon conduction velocity in the model v = v_c = 0.1 m/s (Fig. 2C-D). If we consider instead a lower excitatory coupling *K^E^*(*x, y*) (but still uniform), we observe that the activity that is triggered by the external stimuli in C2 does not propagate with the same intensity in space (Fig. 2E). As a result, the emergence and the spatial structure of the propagating wave are crucially shaped by the average value and the spatial structure of *K^E^*(*x,y*). In the next section we show how it is possible to employ the model in an inverse way, in order to estimate *K^E^*(*x, y*) directly from VSD recordings.

**Figure 2:**
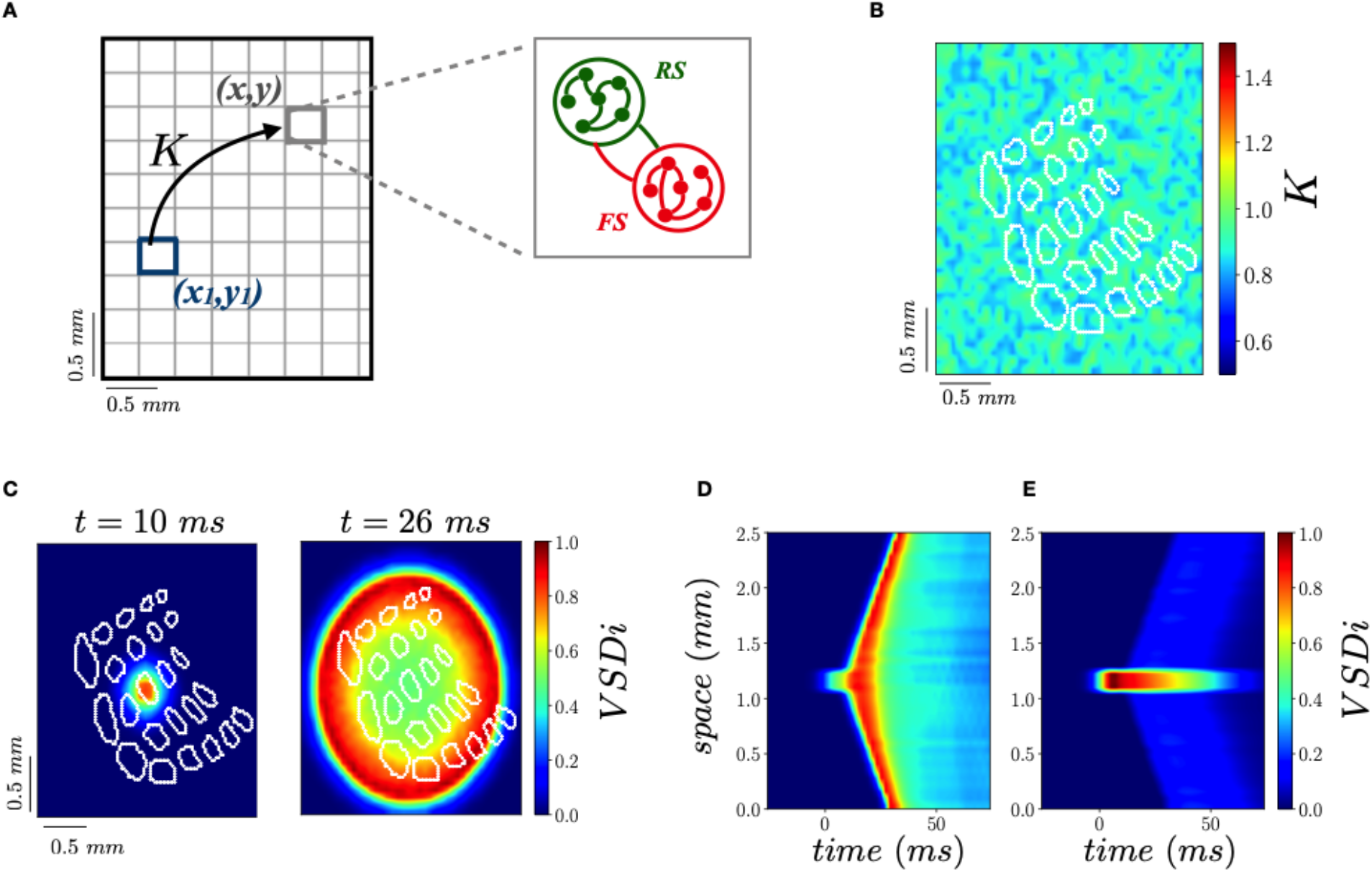
Computational model of S1. **A:** A two dimensional lattice is considered to model activity in S1. Each point at a location (x,y) is modelled as a population of adaptive excitatory Regular Spiking (RS) and non-adaptive Fast Spiking (FS) inhibitory neurons. **B:** The strength of the excitatory incoming synapses (quantal conductance) Q_E_ = Q^0^_E_* K(x,y) is homogeneously distributed around its average Q^0^_E_ (K is extracted from a flat distribution with average <K> = 1 and width ΔK = 0.4). The color plot shows the values of K in the lattice for this flat and homogeneous distribution. **C:** Activity pattern with the distribution as in **B**: a pulse of stimulation in C2 at time t = 0 ms evokes an isotropic propagating wave. **D:** Spatio-temporal activity along the vertical direction passing by C2, same parameters as in **C**. **E:** Spatio-temporal activity along the vertical direction passing by C2 with a uniform *K*, as in D, but with a lower average <*K*> = 0.4 (instead of <*K*> = 1 in panel **B-C-D**).

### Inferring spatial heterogeneity from propagating waves in sensory cortex

We develop here a method to infer the coupling distribution *K^E^*(*x,y*) from the VSD measure of activity in S1. From the measured spatio-temporal VSD pattern, we reconstructed *K^E^*(*x,y*) through a minimization procedure. Starting from an uniform *K^E^*(*x, y*) (we have also used different initial conditions) we search for the distribution *K^E^*(*x,y*) such that the spatio-temporal activity measured experimentally matches the one predicted by the model. We start with a flat uniform distribution (as in Fig. 2B) and we modify *K^E^*(*x,y*) until the relative error between known and predicted VSD activity (integrated in space and time) is smaller than 10 %. More details on the procedure are found in the method section. The inversion procedure could potentially be applied also to other parameters such as *K^I^*(*x,y*). Nevertheless, we focus here on the excitatory coupling keeping uniform *K^I^*(*x,y*) because the response of a population of neurons depends mainly on the ratio between excitation and inhibition (Di Volo et al., 2019). In this view, modifying *K^E^*(*x,y*) and freezing *K^I^*(*x,y*) can be seen as modifying the ratio between excitatory and inhibitory coupling. Nevertheless, in future work we plan to extend the method to reconstruct both the excitatory and inhibitory couplings (see discussion section).

First of all we check the validity of the procedure by considering a well known *K^E^*(*x,y*) with a non-uniform structure (Fig. 3A). We then generate from numerical simulations of the model a synthetic version of VSD data where a propagating wave is originated by a pulse of stimulation in C2 (see methods). These synthetic VSD data are characterized by a spatially non-uniform propagating wave (Fig. 3B). We then employ the inversion procedure from the knowledge of only the synthetic spatio-temporal VSD signal numerically simulated (the spatio-temporal patterns as in Fig. 3B). We observe that the inversion procedure converges to a *K^E^*(*x,y*) very similar to the real one (compare Fig. 3C and Fig. 3A). Moreover, the wave predicted by the model with this reconstructed *K^E^*(*x,y*) is indeed close to the real one (see Fig. 3D vs 3B).

**Figure 3:**
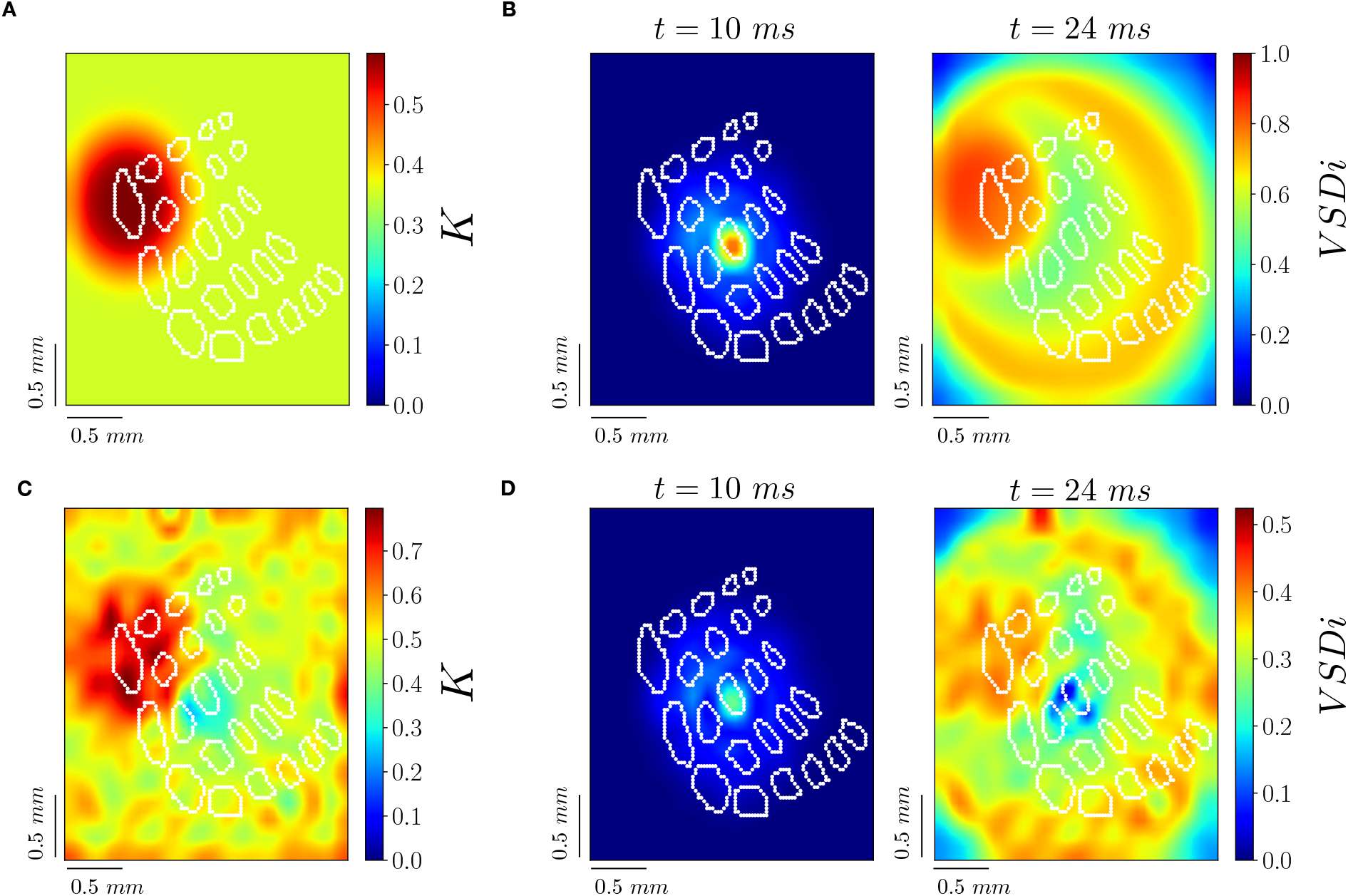
Reconstruction of spatial couplings from synthetic VSD imaging data. **A:** Spatial distribution of couplings K. **B:** Propagating wave emerging in response to a C2 whisker stimulus (at t = 0 ms) in the computational model with couplings as in **A**. **C:** Reconstructed distribution of coupling K with the inversion procedure applied to the VSD activity from **B**. **D:** Propagating wave with the reconstructed distribution of coupling K as in **C**.

Once validated the procedure on synthetic VSD data, we apply the same procedure to the experimental signal (data illustrated in Fig.1). In real data we notice that a wave of activity propagates from the top of the image toward the center at time t = 26 ms (Fig. 4A). This wave is probably originating from the secondary somatosensory cortex S2, as suggested in (Hubatz et al., 2020). As our model does not account for the interaction between S1 and S2, we consider the spatio-temporal signal up to time t = 28 ms and we do not take into account the spatial activity on the top of the image. The inversion procedure converges to a reasonably good match between the spatio-temporal pattern observed experimentally and the one predicted by the model (see Fig. 4A). Notice that with the initial guess (i.e. a flat *K^E^*(*x, y*)), we get the uniform pattern of Fig. 2D. The method is thus able to successfully reproduce the anisotropy of the propagating wave. Nevertheless, we cannot reproduce the high activity observed at the top of the image (see Fig. 4A time = 26 ms), which may be due to an interaction with S2. We recover an heterogeneous and non-symmetric distribution of couplings *K^E^*(*x,y*) (Fig. 5A) with a non-uniform spatial distribution as reported then in Fig. 5B. The coupling K is concentrated in the sensory representation of whiskers in S1. This indicates that the whiskers representation subfield of S1 is a strongly interconnected network with respect to surrounding areas. Nevertheless, a relatively strong coupling appears at the postero-medial side of the image, out of the whiskers representation. This indicates that this region, which lies partly on the anterior extrastriate area (Hovde et al., 2019), is highly connected with the whiskers subfield of S1.

**Figure 4:**
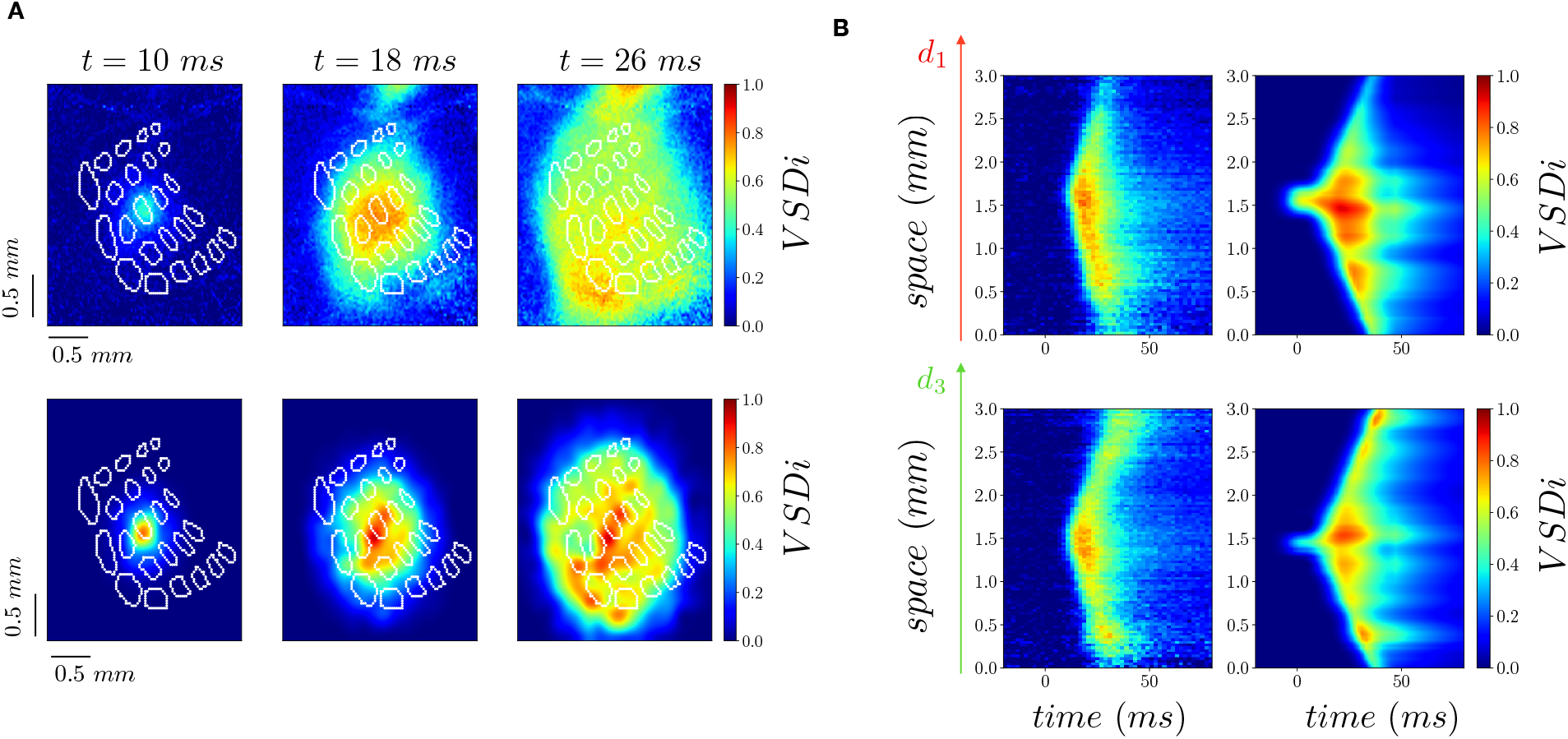
Reconstruction from experimental VSD imaging data. **A:** Time frames of VSD activity evoked by a C2 whisker stimulus (at t = 0 ms) from data (top row) and from the model with reconstructed couplings (bottom row). **B**: Spatio-temporal linescan plots computed from data (left) and the model (right) for two different spatial segments (see Fig. 1**C**).

**Figure 5:**
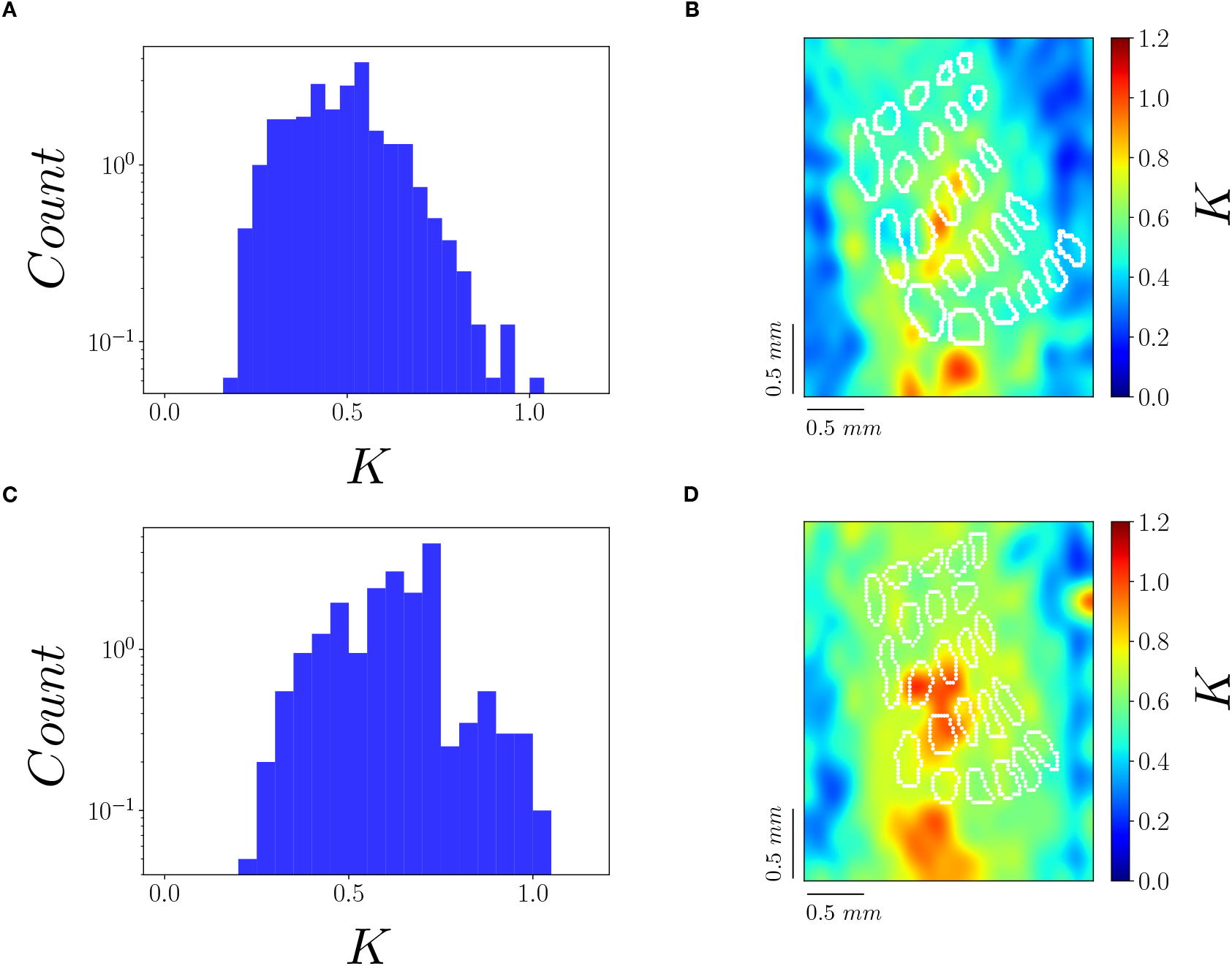
Couplings reconstructed from experimental VSD imaging data. **A:** Histogram of the values of K reconstructed with the inversion method. **B:** The reconstructed spatial distribution of K. **C-D:** As **A-B** for a different mouse.

### Suppressive interaction between propagating waves

In previous sections we have shown that a short stimulus can induce a transition from a lower activity state to a higher activity state that propagates in an anisotropic way in S1 according to the spatially heterogeneous distribution of coupling strengths. The response to an external stimulus delivered at a given spatial location depends on the strength of the input but also on the state of the population of neurons in that specific location. In this network of RS and FS neurons very active states have a lower response to external stimulation. The nonlinearity of population sensitivity to external perturbations is due to the combination of a higher gain of inhibitory with respect to excitatory neurons, combined with conductance based synapses and with spike frequency adaptation (Chemla et al., 2019; Di Volo et al., 2019). Accordingly, in each spatial location, the model predicts a sub-linear response in function of the level of activity. In the spatially extended model here considered, this non-linearity emerges when two propagating waves collide together. Indeed, the response of each subpopulation of neurons to the activity coming from the second wave will be relatively small due to its previous activation from the first wave. Hence, we expect this nonlinearity (more precisely a sub-linearity of response in function of the level of activity) to emerge also in this spatially extended model when two propagating waves collide together. In order to verify this hypothesis, we consider the response evoked by the stimulation of two spatially close whisker representations in S1 (C4 and C2, Fig. 6A-B). Each stimulus provokes a propagating wave that spreads in a few milliseconds to the representation of the other whisker. Consequently, when the two stimuli are presented at a short time interval (Fig. 6C), the waves collide together and the representation of one stimulus overlaps the representation of the other. We measured that the activity evoked by the two stimuli presented sequentially is lower than the linear summation of the responses to the two stimuli presented separately. This can be quantified by considering the difference between the actual response and the linear expectation (normalized by the maximum activity to the 1st stimulus response). As we can see in Fig. 6D, this gives rise to a suppressive wave (negative meaning the actual response is lower than the linear expectation) that propagates from the second stimulated whisker’s representation site to the first one. In order to verify this prediction from our simulations, we have applied this stimulation protocol in experiments and measured VSD activity in S1. More specifically, we have first recorded responses to stimuli applied to the C4 and to the C2 whisker independently, and then measured evoked activity when the C2 whisker is stimulated 10 ms after the C4 whisker, by means of VSD imaging. In Fig.7 we report the results for two different mices. Panel A shows VSD signals imaged in response to a C4 stimulation alone, Panel B shows the response to a C2 stimulation alone, and panel C shows the response to the stimulation of the paired stimulation of the C4 and the C2 whiskers with a delay of 10 ms. In panel D we report the suppressive wave as defined for Fig.6. These results show that, as predicted by the model, the activity of the two colliding waves is lower than the linear prediction from the two waves measured separately.

**Figure 6:**
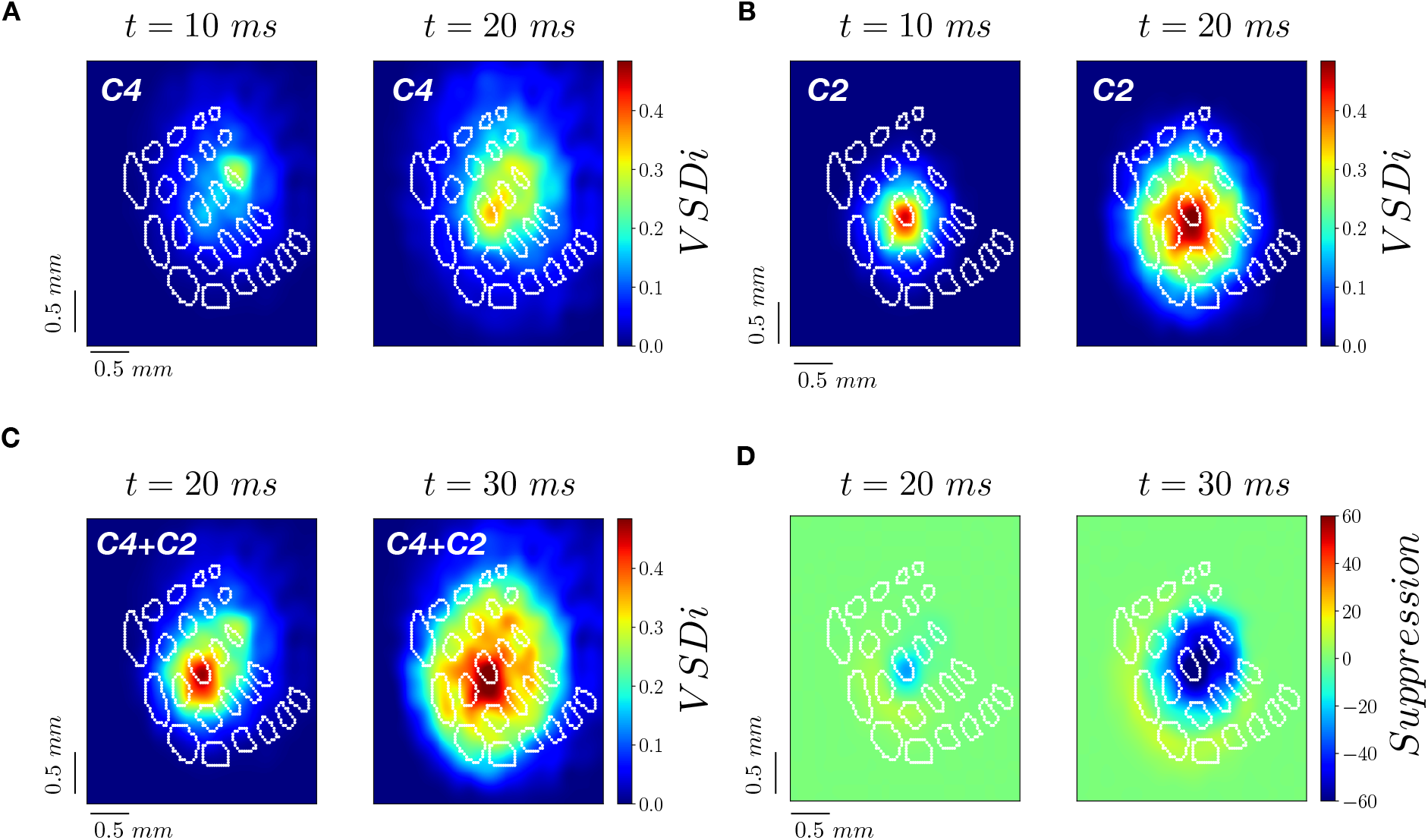
Suppressive interaction between propagating waves in the model. **A:** VSD activity simulated by stimulation of the C4 whisker at t = 0 ms. **B:** VSD activity simulated by stimulation of the C2 whisker at t = 0ms. **C:** VSD activity simulated by the sequential stimulation of the C4 whisker at t = 0 ms and the C2 whisker at t = 10 ms. **D:** Difference between data from experiment as in panel **C**, and the linear prediction from panel **A**+**B** (with proper + 10 ms time shift of data from C2 whisker stimulation), normalized by the maximum of activity in activity of data of panel **A**.

**Figure 7:**
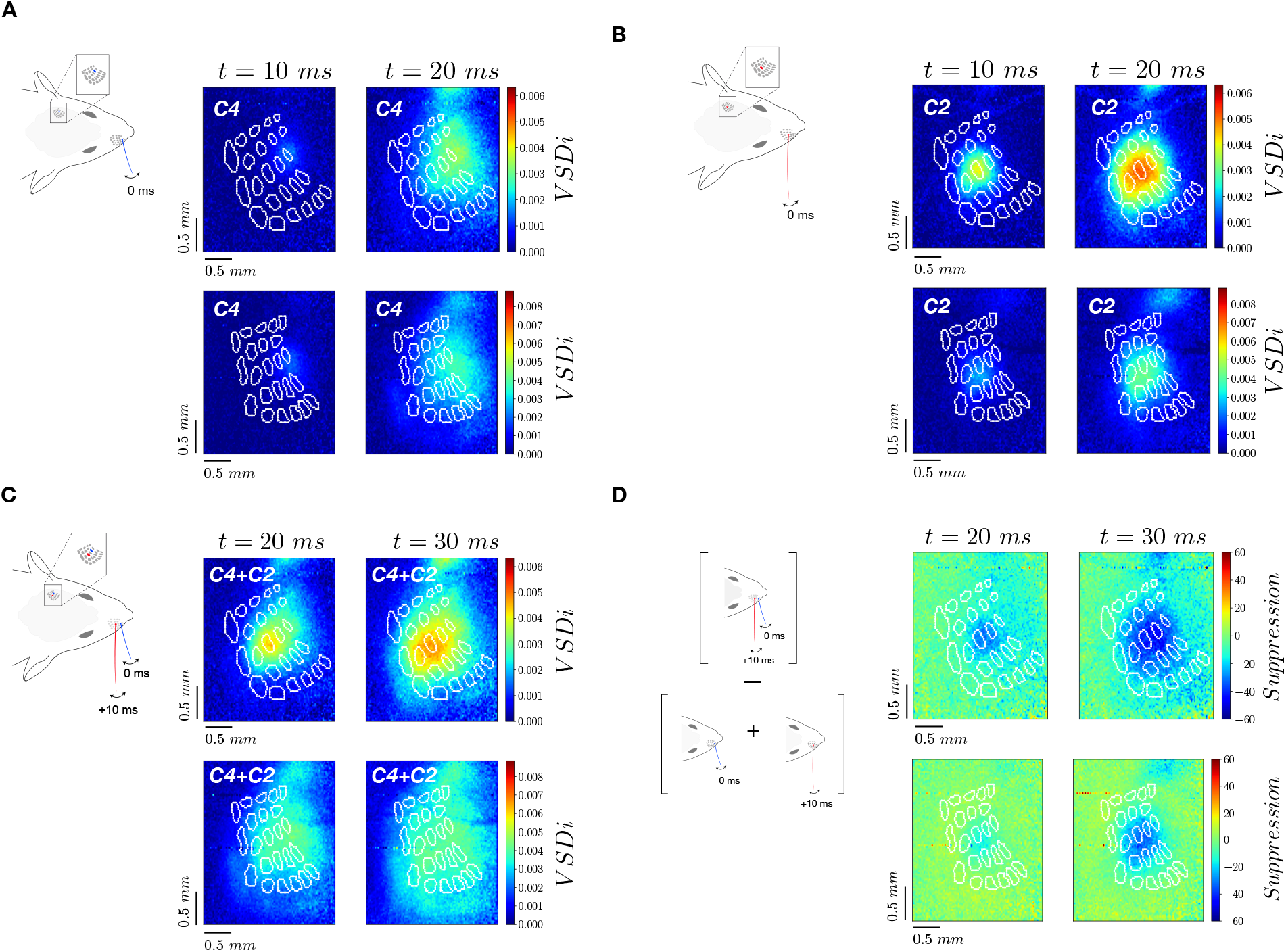
Suppressive interaction between propagating waves in experiment. Different rows in panels **A-D** are extracted from experiments with two different animals. **A:** VSD activity in response to stimulation of the C4 whisker at time t = 0 ms **B:** VSD activity in response to stimulation of the C2 whisker at time t = 0 ms. **C:** VSD activity in response to the sequential stimulation of C4 and C2 whiskers with a delay of 10 ms **D:** Difference between data from experiment as in panel **C**, and the linear prediction from panel **A**+**B** (with proper +10 ms time shift of data from C2 whisker stimulation), normalized by the maximum of activity in data of panel **A**.

## Discussion

In this work we employed VSD imaging and computational modeling to investigate the origin of anisotropic propagating waves of activity in the mouse somatosensory cortex in response to tactile stimulation. We have developed an inversion method that permits to infer the distribution of spatially distributed cellular parameters in the model directly from experimental VSD recordings. Our results show that a spatially heterogeneous distribution of excitatory synaptic strengths is responsible for the emergence of anisotropic pattern of activity measured in S1. This approach can be extended to a large variety of VSD signal recordings, in different regions and animals, permitting to derive different cellular organization directly from large scale imaging measurements in the brain. Finally, we have employed the model to predict, and then to verify experimentally, a suppressive interaction of propagating waves in S1. This suppression emerges from nonlinearities in the sensitivity to perturbations in networks of excitatory and inhibitory neurons.

It is well known that a fraction of more active neurons in S1, giving rise to a skewed distribution of response probabilities across (Barth & Poulet, 2012; O’Connor et al., 2013). A recent modeling work (Moldakarimov et al., 2018) has shown how a subnetwork of highly connected neurons immersed in a larger network of weakly coupled neurons can give rise to propagating waves in S1 where neurons have a heterogeneous spiking activity. In our model, heterogeneity of neural responses emerges as a direct result of our inversion procedure to match VSD experimental recordings (see the skewed distribution in Fig. 5A). Indeed, a neuron with lower strength of incoming excitatory synapses has a lower activation in response to external stimulus. Moreover, our analysis shows that such heterogeneity has a specific structure in space, responsible for the anisotropic wave observed in VSD recordings. Other sources of heterogeneity in our model could explain a different response of neurons, just like neurons resting potential of membrane time constants. While in this work modifications of excitatory synapses strength were enough to match VSD data in response to whisker stimulations, future works should address how heterogeneity in different parameters can affect the structure of propagating waves.

Our model limited its analyses to the early response of S1 after tactile stimulation. As already discussed in the result section, a propagating wave originating outside S1 appears around 20 ms after whisker stimulation and propagates toward whiskers representation in S1. This wave is originating in S2 which receives direct tactile sensory inputs from the thalamus (Carvell & Simons, 1987; El-Boustani et al., 2020; Minamisawa et al., 2018), as well as a monosynaptic drive from S1 through axons travelling in deep cortical layers (Minamisawa et al., 2018). As shown in (Hubatz et al., 2020), S2 comprises a discernable topographic arrangement of individual whisker representations, and responds with a similar intensity to individual whisker stimulation as S1, but with a slight delay in time. It may have a key role in the representation of sensory stimuli (Chen et al., 2013; Kwon et al., 2016; Yamashita et al., 2013; Yamashita & Petersen, 2016; Yang et al., 2015). An extension of this model to account for S2 may shed light on the cellular organization of the whole somatosensory cortex and the interaction between S1 and S2, directly from VSD measurements.

In this work we have employed a recently developed biologically realistic mean field model of the cortex and arranged it in 2D space to compare with VSD recordings of evoked activity. Our approach employs a low dimensional population model keeping track of fundamental biologically realistic ingredients, such as spike frequency adaptation and conductance based synapses. Nevertheless, a different approach has been recently proposed (Newton et al., 2019), where VSD activity is obtained from numerical simulations of a highly detailed model designed directly from cellular information. While we do not have access to cellular resolution with VSD imaging, the advantage of the approach presented here is to keep the model simple and sufficiently realistic. Indeed, the inversion procedure to infer model parameters from VSD data would be computationally impossible if we were ambitioning to model all cellular details. This method succeeded in calibrating model parameters for an optimal match with VSD imaging data and showed that the anisotropic activity of S1 in response to whisker stimulation can be explained from a spatially heterogeneous coupling between neural populations in S1. On the other side, we have shown that the model is able to encode enough realistic features to correctly reproduce the complex nonlinear effects we have investigated, just like the nonlinear interaction between propagating waves in S1. The computational model designed directly from experimental data paves the way to future studies to investigate the activity in S1 in response to more complex simulations. Taking into account spike frequency adaptation, the model gives the opportunity to make predictions of S1 activity in different brain states. In a recent work we have shown that spike frequency adaptation can model neuromodulation, thus explaining differences in cortical dynamics during awake states, sleep and anesthesia (Nghiem et al., 2020). The model here developed will permit to study differences in cortical information processing across brain states.

A nonlinear, suppressive, interaction between propagating waves has been recently reported in the visual cortex of the awake monkey in response to two consecutive and spatially closed visual stimuli (Chemla et al., 2019). We have shown here that this phenomenon appears in a similar protocol also in the somatosensory cortex. The mean field model predicts that this sub-linearity is due to a lower sensitivity to external stimulation of populations of excitatory and inhibitory neurons during high activity states. This prediction can be checked by measuring both the intensity of the suppression and the response of S1 to noisy stimulation, estimating their correlation across different animals and brain states. Moreover the observed suppressive wave, given its robustness across species and brain regions (monkey V1 and mouse S1), is likely to be a general cortical mechanism to discriminate ambiguous stimuli, such as the consecutive stimulation of one whisker followed by the stimulation of another whisker with a time delay Δt (here we used two close whiskers, C2 and C4, and Δt=10ms). Future works should investigate the intensity of the suppressive wave by increasing spatial and temporal distance Δt between the two stimulations, in order to further explore the role of suppressive wave to disambiguate external stimulation.

## Materials and Methods

### Experimental procedure

Experiments were performed in accordance with the French and European (2010/63/UE) legislations relative to the protection of animals used for experimental and other scientific purposes. Experimental procedures were approved by the French Ministry of Education and Research, after consultation with the ethical committee #59 (authorization number: APAFIS#3561-2016010716016314). VSD imaging was performed on six 27–33 days-old C57BL6J mice under isoflurane (induction 3–4%, maintenance 1–1.5%) anesthesia. Paw withdrawal, whisker movement and eye-blink reflexes were suppressed by the anesthesia. A heating blanket maintained the rectally measured body temperature at 37 °C. The respiration of the mice was monitored with a piezoelectric device and the brain state monitored by using two epidural electrodes above the barrel cortex and the frontal cortex ipsilateral to the stimulated whiskers. A metallic fixation post was implanted on the occipital bone with cyanoacrylate glue and dental cement. A ~3 × 3 mm craniotomy was made to expose S1. Extreme care was taken at all times not to damage the cortex, especially during the removal of the dura. The voltage-sensitive dye RH1691 (Optical Imaging Ldt, Israel), dissolved at 1 mg/ml in Ringer’s solution containing (in mM): 135 NaCl, 5 KCl, 5 HEPES, 1.8 CaCl2, 1 MgCl2, was topically applied to the exposed cortex and allowed to diffuse into the cortex over 1 hour. After removal of the unbound dye, the cortex was covered with agarose (0.5–1% in Ringer’s) and a coverslip.

Cortical imaging was performed through a tandem-lens fluorescence microscope (SciMedia Ldt, USA), equipped with a couple of Leica PlanApo objectives, a 100 W halogen lamp gated with an electronic shutter, a 630 nm excitation filter, a 650 nm dichroic mirror, and a long-pass 665 nm emission filter. We set the field of view to 2.5 × 2.5 mm by using a 5x objective on the cortex side, and a 1x objective on the camera side. Images were acquired with a high-speed MiCam Ultima camera (SciMedia Ldt., USA) at 500 Hz. The illumination of the cortical surface started 500 ms before each image acquisition to avoid acquiring signal in the steeper phase of the fluorescence bleaching. Recordings were then of 1 second duration, with 200 ms baseline and 800 ms post stimulation. Variations of the fluorescence signals were initially recorded as variations over the resting light intensity (first acquired frame).

Individual deflections of the right C2, C4 whisker, or paired deflection of the C4 followed by the C2 whisker (with a 10 ms delay) were performed using a multi-whisker stimulator (Jacob et al., 2010) at 0.1 Hz within pseudo randomized sequences containing blank trials (each stimulation being repeated 40 times). Whiskers on the right side were cut to a length of 10 mm and inserted, while keeping their natural angle, in 27G stainless steel tubes attached to piezoelectric benders (Noliac, Denmark), leaving 2 mm between the tip of the tube and the whisker base. Each whisker deflection consisted of a caudal 95 μm displacement (measured at the tip of the tube), a 2 ms rising time, a 2 ms plateau and a 2 ms fall. Specific filters were applied to the voltage commands to prevent mechanical ringing of the stimulators.

Following the experiments mice were perfused with saline followed by paraformaldehyde (4% in 0.1 M phosphate buffer). After an overnight post-fixation in paraformaldehyde, the brains were cut in 100 mm-thick tangential sections that were stained for cytochrome oxidase. Microphotographs of the tangential sections were registered and the barrel maps reconstructed using a method implemented in MATLAB (MathWorks, USA), as previously described (Perronnet et al., 2016). The functional VSD data were aligned with the reconstructed barrel maps by using the superficial blood vessels as anatomical landmarks.

Acquisition and data preprocessing were done using in-house software (Elphy, G. Sadoc, UNIC-CNRS). Subtraction of a pixel by pixel best fit double-exponential from the averaged unstimulated sequence was used to correct for photobleaching.

## Computational model

We consider a two dimensional square lattice. Every node of the lattice represents the network activity of a large population of excitatory Regular Spiking (RS) neurons and inhibitory (FS) fast spiking neurons (Fig. 1b-c).

### Network model

We consider Adaptive Exponential integrate and fire neurons evolving according to the following differential equations:

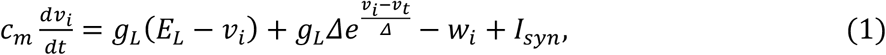

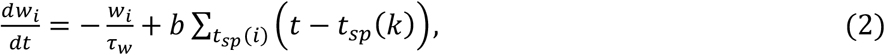

Where *c_m_*=100pF is the membrane capacity, *v_i_* is the voltage of neuron *i* and, whenever *v_i_* >v_th_=-50mV at times t_sp_(k), *v_i_* is reset to its resting value *v_rest_* =-65mV. The leak term has a conductance *g_L_* = 10nS and a reversal *E_L_* = −65mV. The exponential term has a different strength for regular-spiking (RS) and fast-spiking (FS) cells, i.e. *Δ* = 2 mV (*Δ* = 0.5 mV) for excitatory (inhibitory) cells. The variable *w_i_* mimicks the dynamics of spike frequency adaptation. Inhibitory neurons are modeled according to physiological insights as the FS neurons with no adaptation, while the excitatory RS neurons have a lower level of excitability due to the presence of adaptation. Here we consider *b* = 60 pA and *τ_w_* = 500 ms, if not stated otherwise. The synaptic current impinging on the postsynaptic neurons *k, l_syn_*, is modeled as:

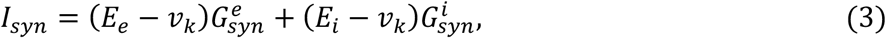

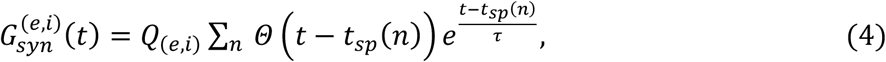

where *Q_e_* (*Q_i_*) is the excitatory (inhibitory) quantal conductance. The variable *τ* = 5 ms is the decay timescale of excitatory and inhibitory synapses, and *Θ* is the Heaviside step function. The summation runs over the over all the pre-synaptic spiking times *t_sp_*(*n*). We set *Q_e_* = 1.5 nS and *Q_i_* = 5 nS. We then consider a random network with p=5% of connectivity and 80% of excitatory neurons. The parameters are chosen according to biological realism for which this model gives rise to asynchronous irregular activity states as observed in the primate visual cortex (Di Volo et al., 2019).

### Spatially extended two dimensional mean field model

The activity of the network is simulated using a mean field model, capable to predict spontaneous activity in both asynchronous and bistable UP-DOWN states dynamics (Di Volo et al., 2019). Connecting several mean field models in space, we obtain the following equations for the spatially extended lattice model:

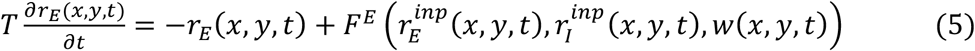

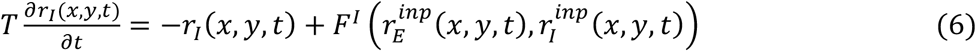

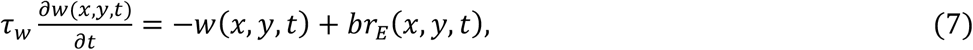

where r_E/I_(x,y) represents the instantaneous firing rate of excitatory/inhibitory neurons in the location (x,y) and F^E/I^ is the transfer function of excitatory/inhibitory neurons. The variables 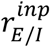 are the net input received by node (x,y) from the rest of the lattice, which can be written as:

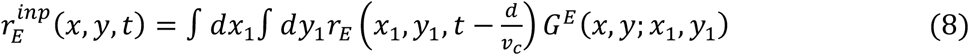

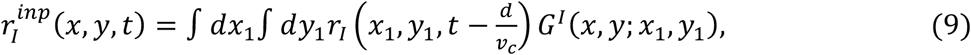

where *d* is the distance between (x,y) and (x_1_, y_1_), *d*^2^ = (*x* – *x*_1_)^2^ + (*y* – *y*_1_)^2^ and *G^E^*(*x, y*; *x*_1_, *y*_1_) is the effective excitatory coupling between the node (x,y) and the node (x_1_, y_1_). The parameter v_c_=0.1mm/ms is the axonal conduction speed, and T = 5 ms is the decay time of population rate. The functions F^E,I^ are the transfer functions of excitatory/inhibitory neurons and are calculated according to a semi-analytical method (Di Volo et al., 2019; Zerlaut et al., 2018) through an expansion in function of the three statistics of neurons voltage, i.e. its average *μ_V_*, its standard deviation *σ_V_* and its autocorrelation time *τ_V_*:

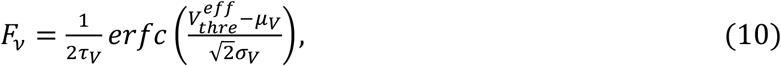

where *erfc* is the Gauss error function, 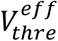 is an *effective* or *phenomenological threshold*. This threshold is expressed as a first order expansion with some fitting coefficients in function of (*μ_V_, σ_V_, τ_V_*), which are calculated from shot-noise theory (Kuhn et al., 2004). Introducing the following quantities:

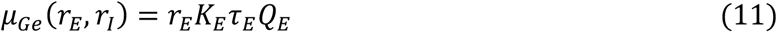

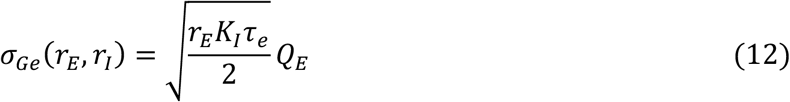

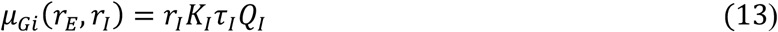

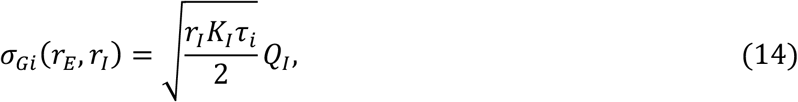

where K_E/I_ is the amount of incoming synapses related to pre-synaptic excitatory/inhibitory neurons (we consider a network of N=10000 neurons inside each node of the ring), we obtain the following equations for the voltage moments:

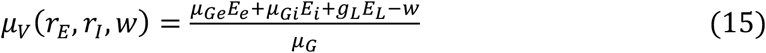

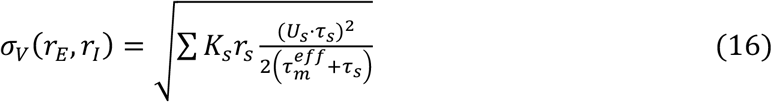

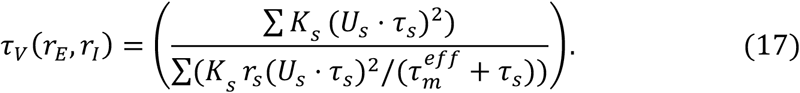

Details on fitting procedure and the comparison between mean field predictions and network simulations are reported in (Di Volo et al., 2019).

### Spatial coupling

The function G(x,y,x_1_,y_1_) represents the effective directional coupling from site (x1,y1) to site (x,y). Notice that, in a mean field description as the present one, this coupling can be interpreted also as the average amount of incoming connection that a neuron in (x,y) receives from a neuron in (x_1_,y_1_). We assume this coupling to be decreasing with the distance *d* between (x_1_,y_1_) and (x,y), and proportional to the average amount of incoming inputs received by a neuron in (x,y), K(x,y):

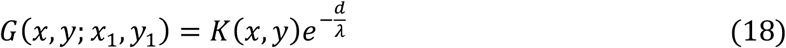

where λ is a scale determining the decay and K(x,y) can be interpreted as an effective coupling of (x,y) with the rest of the lattice. In this work we have fixed λ = 0.8 mm (notice that the lattice we consider has an edge L = 2.5 mm). We have also verified that small modification of the value of λ did not affect the results of our analyses.

## Inversion procedure

We search for the coupling function K(x,y), such that the spatio-temporal pattern measured experimentally matches the one predicted by the model. We consider an experimental setup where a propagating wave is activated in C4. Accordingly, all neurons in C4 receive a fast and spatially local external excitatory input that activates the propagating wave. The input is an excitatory poissonian train of spikes of duration T_inp_ = 2 ms and amplitude A_inp_ = 10 Hz. We then consider a coarse grained version of the spatial model with M = 20 pixels in each direction. The function K(x,y) is then a MXM matrix. For a specific matrix K we simulate the model and obtain a time varying 2D matrix VSDi_model_ to be compared with VSDi_exp_. We introduce the distance D such that:

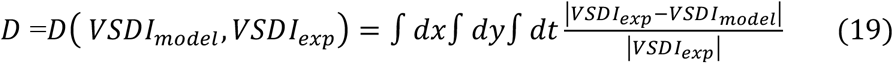

The inversion procedure works as follows: we start with an homogeneous matrix K_0_(x,y) as initial conditions and, at each step n=1..N of the procedure, we pick randomly a location (x_c_, y_c_) and increase or decrease the value of the matrix of a quantity ε. We thus obtaine the matrix K_n_(x,y) such that K_n_(x_c_, y_c_)=K_n-1_(x_c_, y_c_) +ε. We then simulate the 2D mean field model with K_n_ obtaining VSDi_model_(n) and the new distance D_n_= D(VSDi_model_(n), VSDi_exp_(n)). If the distance d_n_<d_n-1_ we accept the modification to K, if not we set K_n_ = K_n-1_. The procedure continues up to when we reach a satisfactory (smaller then 1) distance d_N_. In the inversion procedure reported here we have chosen ε=0.1, and employed N around 5000 up to a distance of D_N_ arounds 0.1. The time window in which we perform the inversion procedure is Δt = 20ms.

## Acknowledgments

The authors acknowledge A. Destexhe for extremely useful discussion on the manuscript, as well as D. Shulz for enlightening discussions and comments/feedbacks on the manuscript. MDV has received partial support by the French Governement via the ANR Project ERMUNDY (Grant No. 18-CE37-0014-03). Experimental assistance was provided by Aurélie Daret and Guillaume Hucher. Experimental work was supported by Equipe Fondation de la Recherche Médicale (FRM) DEQ.20170336761, the Lidex Neuro-Saclay (BRAINSCOPES, iCode), the Agence Nationale pour la Recherche (SensoryProcessing, Neurowhisk, Expect) and the CNRS.

